# MiR-182-5p regulates Nogo-A expression and promotes neurite outgrowth of hippocampal neurons *in vitro*

**DOI:** 10.1101/2022.03.03.482803

**Authors:** Altea Soto, Manuel Nieto-Díaz, David Reigada, Teresa Muñoz-Galdeano, M. Asunción Barreda-Manso, Rodrigo M. Maza

## Abstract

Nogo-A protein is a key myelin-associated inhibitor for axonal growth, regeneration, and plasticity in the central nervous system (CNS). Regulation of the Nogo-A/NgR1 pathway facilitates functional recovery and neural repair after spinal cord trauma and ischemic stroke. MicroRNAs are described as effective tools for the regulation of important processes in CNS such as neuronal differentiation, neuritogenesis, and plasticity. Our results showed that miR-182-5p mimic specifically downregulates the expression of the luciferase reporter gene fused to the mouse Nogo-A 3’UTR, and Nogo-A protein expression in Neuro-2a and C6 cells. Finally, we observed that when rat primary hippocampal neurons are co-cultured with C6 cells transfected with miR-182-5p mimic, there is a promotion of the outgrowth of neuronal neurites in length. From all these data we suggest that miR-182-5p may be a potential therapeutic tool for the promotion of axonal regeneration in different diseases of the CNS.

**Graphical abstract:** 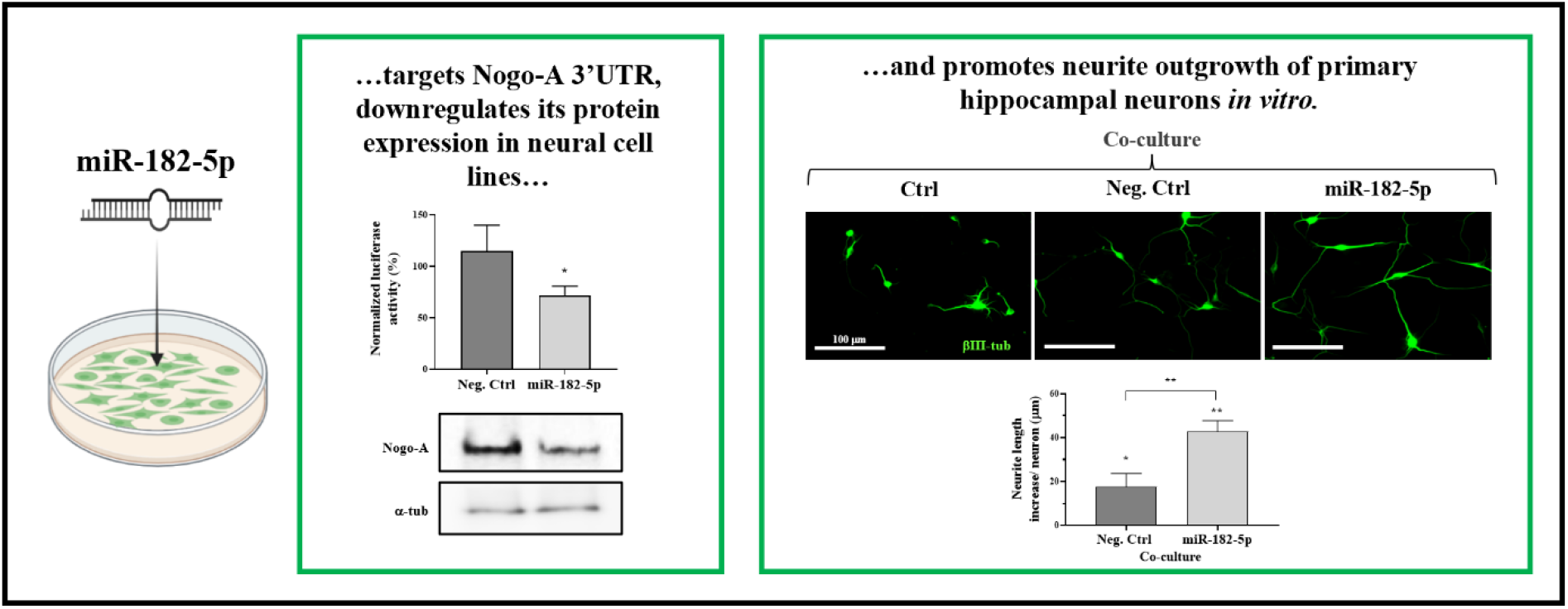

**Highlights:**

- Bioinformatics analyses show that miR-182-5p targets Nogo-A 3’UTR.
- MiR-182-5p downregulates Nogo-A protein expression in murine cell lines.
- MiR-182-5p promotes neurite outgrowth of rat primary hippocampal neurons *in vitro*.
- MiR-182-5p is suggested as a potential therapeutic tool for the promotion of axonal regeneration in different pathologies/diseases of the central nervous system.

## 1. Introduction

During postnatal development, central nervous system (CNS) neurons lose their ability to regenerate in part due to the presence of myelin-derived inhibitors of axonal outgrowth and neuroregeneration, such as MAG, OMGp, and Nogo-A (Montani *et al.,* 2009). Nogo gene (RTN4) belongs to the family of reticulon encoding genes that produces three major protein variants, RTN4A (Nogo-A), RTN4B (Nogo-B), and RTN4C (Nogo-C) by alternative splicing, promoter usage and alternative polyadenylation (Oertle *et al.,* 2003). All of them share a common 188 amino acid C-terminal membrane-spanning domain, known as a reticulon homology domain (RHD domain), consisting of two hydrophobic regions flanking a hydrophilic loop (Nogo-66), which is followed by the short C-terminal tail. Nogo-A is the largest of the Nogo isoforms with two known neurite growth inhibitory domains including amino-Nogo (Nogo-A-Δ20) at the N-terminus and the extracellular loop Nogo-66 (Schwab *et al.,* 2010).

Nogo function depends on the tissular expression of each isoform (Oertle and Schwab, 2003). Nogo-A has been studied extensively in the CNS and is responsible for the inhibition of neurite outgrowth acting as a myelin-associated inhibitor of axon regeneration. Nogo-A is mostly expressed in spinal motor, DRG, sympathetic, hippocampal, and Purkinje neurons as well as oligodendrocytes. In the spinal cord, Nogo-A is mainly expressed in grey matter, especially by large motor neurons of the ventral horns (Josephson *et al.,* 2001; Huber *et al.,* 2002; Hunt *et al.,* 2003). The functions carried out by Nogo-A in neurons and oligodendrocytes appear to be quite different. While neuronal Nogo-A seems to act as a direct local restrictor of synaptic and dendritic plasticity, oligodendrocytic Nogo-A may act as an inhibitor of axonal growth through binding to its receptor (Nogo receptor, NgR1) located on neurons. This binding transduces the inhibitory signal to the cell interior via transmembrane co-receptors LINGO-1 and p75NTR or TROY by which the small GTPase RhoA is activated (Montani *et al.,* 2009; Schwab *et al.,* 2010; Fournier *et al.*, 2001; Aloy *et al.*, 2006; Zemmar *et al.*, 2018). Multiple studies have demonstrated the efficacy of targeting Nogo-A/NgR1 pathway for functional recovery and neural repair after spinal cord trauma, ischemic stroke, optic nerve injury, and models of multiple sclerosis (see review by Schwab and Strittmatter, 2014 and references therein).

MicroRNAs (miRNAs) are an abundant class of small non-coding RNAs that operate as epigenetic modulators of gene expression in physiology but also in pathophysiological processes. They are involved in post-transcriptional gene silencing by base-pairing to their target mRNAs of protein-coding genes resulting in reduced translation of the protein by mRNA repression or degradation (Bartel *et al.*, 2009; Moore *et al.*, 2015). The flexibility and efficiency of miRNA function provide both spatial and temporal gene regulatory capacities that are essential for establishing neural networks. The expression of miRNAs is ubiquitous in neural tissues, and many of them regulate neuronal differentiation, neuritogenesis, excitation, synaptogenesis, and plasticity (Kocerha *et al.*, 2009; Coolen and Bally-Cuif, 2009; Moore *et al.*, 2015). There is a close relationship between miRNAs and intrinsic determinants of axonal regeneration. Several miRNAs have been proven to regulate the expression levels of targets involved in neurite outgrowth and axonal regeneration after CNS injury. For example, miR-431, one of nerve injury-induced miRNAs that stimulates regenerative axon growth by silencing Kremen1, an antagonist of Wnt/beta-catenin signaling (Wu *et al.,* 2013) or miR-133b which has also been shown to be promoting neurite outgrowth in primary cortical neurons and PC12 cells by targeting RhoA (Lu *et al.,* 2015). Other examples are miR-124, a well-conserved brain-specific miRNA that promotes neurite outgrowth of M17 cells by targeting ROCK1 GTPase (Gu *et al.*, 2014), or miR-222 that regulates neurite outgrowth from DRG neurons by targeting PTEN (Zhou *et al.*, 2012).

However, a functional validated miRNA that regulates the expression of Nogo-A has not been yet described. Even though the reticulon (RTN) family isoforms mature through splicing and alternative polyadenylation processes, Nogo-A gene shares the highly conserved carboxy-terminal reticulon domain and 3’UTR. We thus seek functional validated miRNAs capable of inhibiting the activity of the Nogo family members and only miR-182-5p was found to directly targets Nogo-C 3’UTR and decrease Nogo-C protein levels in cardiomyocytes cells (Jia *et al.,* 2016). MiR-182-5p is a member of the miR-183 family located on chromosome 7q31-34 and is described as an oncogenic miRNA due to its capacity to enhance cancer cell proliferation, survival, tumorigenesis, and drug resistance (Wang *et al.,* 2015; Wei *et al.,* 2015). While miR-182-5p roles are well known in cancer, little is known about its function in the CNS under normal and pathophysiological conditions. Wang and cols. demonstrated that miR-182-5p promotes axonal growth and regulates neurite outgrowth via the PTEN/AKT pathway in cortical neurons (Wang *et al.,* 2017). Moreover, miR-182-5p is the most abundant miRNA in retinal ganglion cell axons, where regulates Slit2-mediated axon guidance *in vitro* and *in vivo* (Bellon *et al.,* 2017). Furthermore, Yu and cols. showed that miR-182-5p inhibits Schwann cell proliferation and migration by targeting FGF9 and NTM, in sciatic nerve injury (Yu *et al.,* 2012).

Since all Nogo protein variants share the conserved carboxy-terminal reticulon domain and 3’UTR, we hypothesize that miR-182-5p could also regulate Nogo-A expression. In the present study, we perform a bioinformatic and validation characterization of the miR-182-5p site in the Nogo-A 3’UTR to demonstrate for the first time that miR-182-5p downregulates Nogo-A protein expression in Neuro-2a and C6 cells and promotes neurite outgrowth of rat primary hippocampal neurons *in vitro*.

## 2. Materials and Methods

### 2.1. Bioinformatics and data mining

To identify miR-182-5p response elements in mouse messenger RNAs (mRNA), an *in silico* screening approach was used. This approach combines computational tools that employ existing databases and prediction algorithms, and data mining for gene expression data analysis. The four prediction tools used were: miRMap (https://mirmap.ezlab.org/, last accession in Nov. 2021), miRanda 3.3a (http://www.microrna.org, Nov. 2021), TargetScan 8.0 (http://www.targetscan.org, Nov. 2021), and miRWalk 3.0 (http://mirwalk.umm.uni-heidelberg.de/, Nov. 2021).

MiR-182-5p binding site accessibility on the human Nogo-A 3’UTR (3’UTR-Nogo-A) was analyzed using STarMir tool (Rennie *et al.,* 2014), an implementation of logistic prediction models developed with miRNA binding data from crosslinking immunoprecipitation (CLIP) studies. In the STarMir web (https://sfold.wadsworth.org/cgi-bin/starmirtest2.pl, Nov. 2021), we input hsa-miR-182-5p into the option of “microRNA sequence(s), microRNA ID(s)”. The human 3’UTR-Nogo-A sequence was input into the option of “single target sequence, manual sequence entry”. By choosing “V-CLIP based model (human)”, “Human (homo sapiens)”, and “3’UTR”, a set of parameters (described in http://sfold.wadsworth.org/data/STarMir_manual.pdf) would be displayed in the output window and ready for further analysis.

### 2.2. Cell lines and cultures

Neuro-2a mouse neuroblastoma cells (cat#: CCL-131, ATCC; RRID#CVCL_0470) and C6 rat brain glioma cells (cat#: CCL-107, ATCC; RRID##CVCL_0194) were cultured in Dulbecco’s modified Eagle’s medium (DMEM; Gibco) supplemented with 10% fetal bovine serum (FBS; Gibco), 1% penicillin/streptomycin (Gibco) and 1% glutaMAX (Gibco). SH-SY5Y human neuroblastoma cells (cat#: CRL-2266, ATCC; RRID#CVCL_0019) were grown in a 1:1 combination of Minimum Essential Medium (MEM; Gibco) and Ham’s F-12 nutrient mixture (Gibco) supplemented with 10% FBS, 1% penicillin/streptomycin, 1 mM sodium pyruvate (Gibco) and non-essential amino acids (Gibco). Cells were cultured at 37°C in a humidified incubator containing 5% CO2.

Primary hippocampal neurons were obtained from 17-18 days old (E17-18) Wistar rat embryos. Briefly, after dissection, hippocampi were subjected to enzymatic digestion in Hanks′ Balanced Salt Solution (HBSS) medium without calcium and magnesium (Hyclone, GE Healthcare) supplemented with Trypsin (1x; Thermo Fisher Scientific) and DNAse (20 mg/ml; Roche) for 15 minutes at 37°C. Trypsin and DNAse were washed out with HBSS with calcium and magnesium (Hyclone) and mechanically disrupted by passing the tissue sample several times through a glass pipette in Neurobasal Medium (Gibco), supplemented with 2% B-27 supplement (Gibco), 1% GlutaMAX (Gibco), and 1% penicillin/streptomycin. Neurons were cultured on C6 cells at 37°C in a humidified incubator containing 5% CO2.

### 2.3. RNA extraction and quantitative real time-PCR (RT-qPCR)

Total RNA was isolated from Neuro-2a, C6, and SH-SY5Y cells employing miRNeasy Kit (Qiagen) and was analyzed with a NanoDrop ND-1000 spectrophotometer (Thermo Fisher Scientific) to determine its concentration and purity (260/280 and 260/230 ratios). To determine cellular miR-182-5p expression, 10 ng of total RNA was reverse-transcribed and amplified using TaqMan miRNA gene expression assay (TaqMan® MicroRNA assay #002284, Applied Biosystems) following manufacturer’s protocols. The U6 small nuclear RNA served as an internal control (TaqMan® MicroRNA assay #001973, Applied Biosystems). The abundance of the microRNA was measured in a thermocycler ABI Prism 7900 fast (Applied Biosystems) applying 40 cycles of a two-step protocol: 15 seconds at 95°C plus 1 minute at 60°C; ΔCt value was defined as the difference between the cycle threshold of amplification kinetics (Ct) of the target miRNA and its respective U6 loading control (Livak and Schmittgen, 2001).

### 2.4. Vector construction

The sequence of the mouse Nogo-A 3’UTR (NM_194054.3) (3’UTR-Nogo-A) containing the predicted binding site for mmu-miR-182-5p (nt 4272-4279) wild-type (wt) was obtained by amplification by PCR from mouse brain total RNA preparation (Table 1). The 3’UTR-Nogo-A-wt product was subcloned into both the pGEM-T-easy plasmid (Promega) and the pBKS plasmid (pBluescript, Stratagene). The 3’UTR-Nogo-A-wt sequence was validated by DNA sequencing (T7p and SP6), and inserted into the pmiRGLO Dual-luciferase miRNA Target Expression Vector (Promega, http://www.addgene.org/vector-database/8236/) between the SacI and XbaI restriction sites (pmiRGLO-3’UTR-Nogo-A-wt) using the FastDigest restriction enzymes (Thermo Fisher Scientific). The miR-182-5p mutant site of 3’UTR-Nogo-A (3’UTR-Nogo-A-mut) was obtained by PCR using the 3’UTR-Nogo-A-wt subcloned into pBKS plasmid as template, specific primers (Table 1), PfuI polymerase (Thermo Fisher Scientific), following by the DpnI endonuclease restriction digest (FastDigest, Thermo Fisher Scientific) and a transformation in *E.coli* supercompetent cells (Thermo Fisher Scientific). The 3’UTR-Nogo-A-mut fragment was inserted into the pmiRGLO plasmid between the SacI and SalI restriction sites (pmiRGLO-3’UTR-Nogo-A-mut). Finally, we confirmed the sequence of both pmiRGLO 3’UTR-Nogo-A constructs by DNA sequencing using a specific forward 3′ end luciferase primer.

**Table 1.**
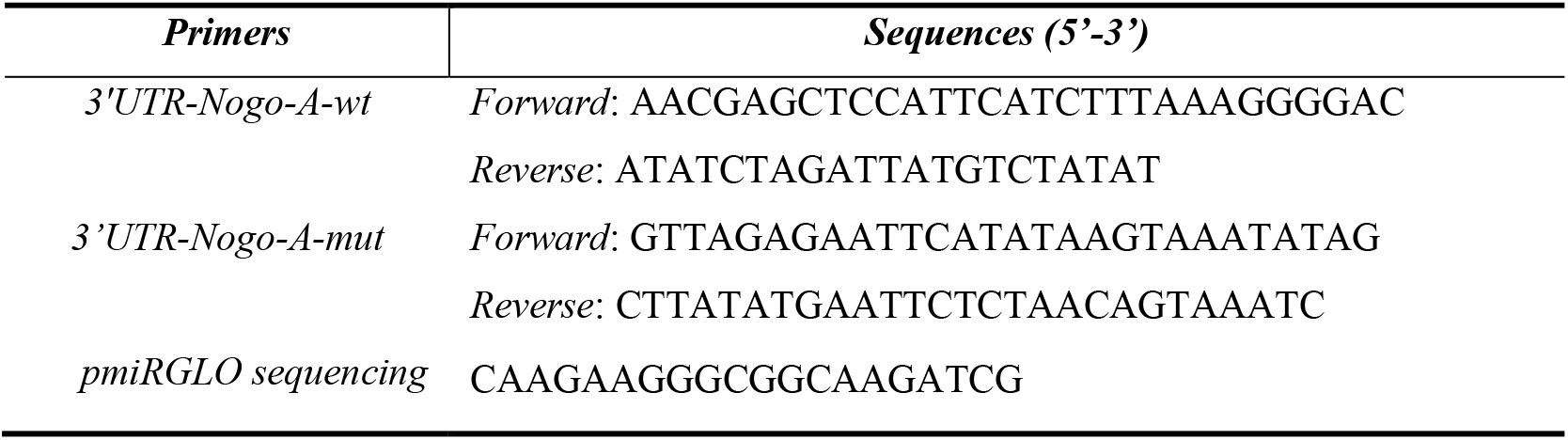
Primers used for subcloning of 3′UTR-Nogo-A-wt and 3’UTR-Nogo-A-mut fragments, and for DNA sequencing.

### 2.5. Dual-luciferase reporter assay

To validate the targeting of miR-182-5p mimic on mouse 3’UTR-Nogo-A, Neuro-2a cells were cultured to 70% confluence in white 96-well plates. Then, cells were co-transfected using DharmaFECT Duo Transfection Reagent (Dharmacon™) with either i) 50 nM of miR-182-5p mimic (miRIDIAN; cat#: C-320575-01-0005, Dharmacon™) or 50 nM cel-miR-67 negative control mimic (Neg. Ctrl mimic) (miRIDIAN microRNA mimic negative control#1; cat#: CN-001000-01; Dharmacon™) and ii) 200 ng/well of pmiRGLO-3’UTR-Nogo-A-wt, pmiRGLO-3’UTR-Nogo-A-mut or empty pmiRGLO (without subcloned 3′UTR) as endogenous regulation control. 24 hours later, the plasmid gene expression under the regulation of both Nogo-A 3’UTRs was evaluated by measuring firefly and Renilla luciferase activities using the Dual-GLO luciferase assay system (Promega) in an Infinite M200 plate reader (Tecan) according to the manufacturer’s protocol. Firefly emission data was normalized to Renilla load control levels and expressed as firefly/Renilla ratio. All experiments were performed in triplicates in, at least, five independent experiments.

### 2.6. Immunoblot assay

To evaluate the endogenous expression levels of Nogo-A protein in different neural cell lines, Neuro-2a, C6 and SH-SY5Y cells were cultured to 70% confluence in 24-well plates. To evaluate the regulation of miR-182-5p mimic on Nogo-A expression, Neuro-2a and C6 cell cultures were transfected using DharmaFECT 4.0 Transfection Reagent (Dharmacon™) with either 50 nM of miR-182-5p or Neg. Ctrl mimics. 24 hours later, cells were lysed with radioimmnunoprecipitation assay lysis buffer (RIPA, Sigma-Aldrich) containing a complete EDTA-free protease inhibitor cocktail (Roche) during 30 minutes at 4°C and cleared by centrifugation (12000xg/10 min/4°C). Protein concentration was determined by the bicinchoninic acid method (BCA protein assay kit, Thermo Fisher Scientific). 25 μg of protein were resolved by 8% dodecyl sulfate-polyacrylamide gel electrophoresis (SDS-PAGE) and then electrophoretically transferred to a 0.45 μm polyvinylidene difluoride membrane (PVDF; Immobilon, Merck Millipore). Membranes were blocked in blocking buffer (5% non-fat milk or 5% FBS diluted in TBS-T buffer (0.05% Tween-20 (Sigma-Aldrich) in tris-buffered saline (TBS)) during 30 minutes at 37°C and immunoblotted with a rabbit polyclonal antibody against Nogo-A (1:100; Santa Cruz Biotechnology, cat#: sc-25660, RRID:AB_2285559) overnight at 4°C. Mouse monoclonal antibody against α-tubulin (1:10000; Sigma-Aldrich, cat#: T6074, RRID:AB_477582) was used as a loading control. Then, membranes were incubated with a horseradish peroxidase (HRP)-conjugated goat anti-rabbit secondary antibody (1:1000; Thermo Fisher Scientific, RRID:AB_228341) or a HRP-conjugated goat anti-mouse secondary antibody (1:1000; Thermo Fisher Scientific, RRID:AB_228307) for 1 hour at room temperature (RT). Finally, protein bands were visualized using the SuperSignal West Pico Chemiluminescent detection system (Pierce, Thermo Fisher Scientific) and measured using ImageScanner III and ImageJ software (National Institutes of Health, Bethesda, MD).

### 2.7. Neurite outgrowth assay

To evaluate the effect of Nogo-A regulation by miR-182-5p on neurite outgrowth of primary hippocampal neurons, different co-cultures of C6 cells (non-transfected cells; Neg. Ctrl mimic transfected cells; miR-182-5p mimic transfected cells) and rat primary hippocampal neurons were performed. C6 cells were cultured to 70% confluence in poly-L-lysine (Sigma-Aldrich) pre-coated 24-well plates. 24 hours later, cells were transfected using DharmaFECT 4.0 Transfection Reagent (Dharmacon™) with either miR-182-5p or Neg. Ctrl mimics. 24 hours later, the medium was changed by supplemented Neurobasal Medium (described above), and 5,000 primary hippocampal neurons were cultured on C6 cells for 24 hours more. The co-cultures were fixed with 4% paraformaldehyde for 30 minutes at RT and washed with 1x Phosphate-buffered Saline (PBS). The cells were permeabilized and blocked with 0.2% Triton X-100 and 3% Bovine serum albumin protein (BSA) respectively, in PBS for 30 minutes at RT. Then, cells were immunostained with a neuronal-specific marker, mouse anti-β−III tubulin isoform antibody (1:500, Millipore, cat#MAB1637, RRID:AB_2210524), for 2 hours at RT and followed by a goat anti-mouse Alexa Fluor 488 conjugated secondary antibody (1:500, Molecular Probes, cat# A-11029, RRID:AB_138404) for another 2 hours at RT. After three washes with PBS, cells nuclei were stained with DAPI (4′,6-diamidino-2-fenilindol, 1:10000, Merck, cat#D9542) for 5 minutes at RT and mounted in Lab Vision™ PermaFluor™ Aqueous Mounting Medium (Thermo Fisher Scientific, cat#TA-030-FM). Co-cultures were imaged in an epifluorescence microscope (DM5000B, Leica Microsystem GmbH), Wetzlar, Germany) with an 40x objective and analyzed using ImageJ software.

Neurite outgrowth lengths were assessed using the method described by Rønn and cols. (Rønn *et al.,* 2000). Briefly, the absolute neurite length (L) per neuron was estimated by counting the number of intersections (I) between neurites and test lines of a grid superimposed on the co-culture images and the the equation L=(πd/2)I, being d the vertical distance between the test lines of the grid. The neurite length increase per neuron was calculated using the Ctrl co-culture (with non-transfected C6 cells) as reference, that is, subtracting the mean neurite length per neuron of the Ctrl co-culture to the mean neurite length per neuron of the mimics transfected co-cultures. C6 cells and hippocampal neurons density were analyzed calculating the total number of both cells per mm^2^ in the different co-cultures (a total of nine images of 0.27 mm2 per condition were analyzed)

### 2.8. Statistical analysis

Statistical significance of the transfection effects was tested using paired Student’s t-test or two-way ANOVA with Tukey’s multiple comparisons post-hoc tests. Data are expressed as mean ± SEM as indicated in figure legends. Statistical analyses and graphic design were conducted using GraphPad Prism version 8.0.0 (GraphPad Software). Differences were considered statistically significant when the p-value was below 0.05.

## 3. Results

### 3.1. MiR-182-5p is predicted to regulate mouse, rat and human Nogo-A 3’UTRs

A bioinformatics-based prediction of the potential targets of miR-182-5p in mouse mRNAs was performed. Since the various available programs can yield rather different predictions, we combined miRmap, miRanda 3.3a, TargetScan 8.0 and miRWalk 3.0-programs to search for mouse miR-182-5p gene targets (Fig. 1A). MiRmap (red) identifies a total of 5020 genes as predicted targets of miR-182-5p, whereas miRanda 3.3a (green), TargetScan 8.0 (blue) and miRWalk 3.0 (yellow) list 1248, 1043 and 10519 genes, respectively. A total of common 191 genes are identified by the four prediction programs, including Rtn4 (Nogo gene).

**Figure 1.**
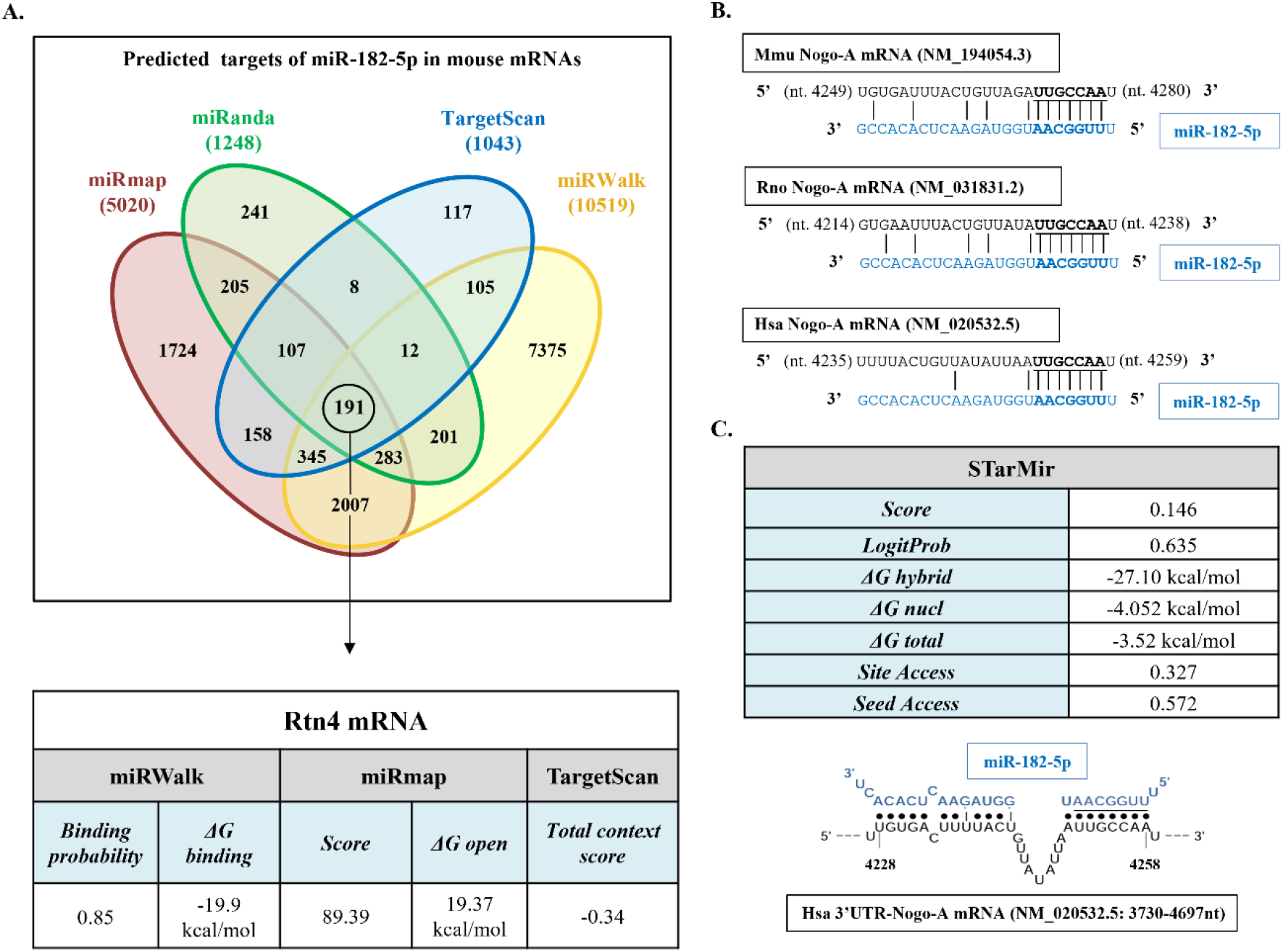
Bioinformatics analyses. **A)** Description and main data of the process of identification of mouse Rtn4 gene (Nogo-A) as miR-182-5p predicted target by miRmap, miRanda 3.3a, TargetScan 8.0 and miRWalk 3.0. The table shows the algorithms prediction scores, binding probability, free energy gained by transitioning from the state in which the miRNA and the target are unbound (ΔG open) (kcal/mol) and the state in which the miRNA binds its target (ΔG binding), according to each algorithm. **B)** Alignment of the seed region of miR-182-5p with the Nogo-A 3’UTR in mouse (Mmu), rat (Rno), and human (Hsa) mRNAs. MiR-182-5p sequence appears in blue type and miRNA seed regions appear in bold type. **C)** Main data of the target site accessibility and hybrid diagram seed site of miR-182-5p on human Nogo-A 3’UTR by STarMir tool. MiR-182-5p sequence appears in blue type and miRNA seed region appears underlined.

According to the employed prediction programs, the Nogo-A 3’UTR of all species has one binding site for miR-182-5p. Alignment of this putative site in rat, mouse, and human sequences demonstrates the evolutionary conservation of Nogo-A site among mammalian species (Fig. 1B). Since the three Nogo protein variants share the same 3’UTR region, we observe that this site matches with the already validated miR-182-5p site in the Nogo-C 3′UTR (Jia *et al.,* 2016). Analyses of target site accessibility of the mRNA secondary structure by STarMir tool further supports miR-182-5p targeting on the human Nogo-A 3’UTR. The logistic probability (*LogitProb*) of the Nogo-A 3’UTR site being a miR-182-5p binding site is 0.635 (Fig.1C). In general, a *LogitProb* of 0.5 indicates a fairly good chance of miRNA binding (Rennie *et al.,* 2014). The hybrid diagram seed site (Fig.1C) showed that miR-182-5p has a canonical site at the human Nogo-A 3’UTR with a hybridization energy (ΔGhybrid) of −27.10 kcal/mol, and therefore, thermodynamically the interaction between miR-182-5p with its respective seed region in the human Nogo-A 3’UTR is stable.

Taken together, the bioinformatics approach agrees that miR-182-5p has a potential site in the sequence of Nogo-A 3’UTR in the three species studied, and therefore miR-182-5p can play a biologically relevant role in regulating their expression.

### 3.2. Nogo-A and miR-182-5p expression in neural cell lines

To validate the functional miRNA/mRNA target interaction, the endogenous expression levels of miR-182-5p and Nogo-A protein in different neural cell lines were compared (Mouse: Neuro-2a; Rat: C6; Human: SH-SY5Y). Data suggest that Neuro-2a and C6 cell lines are appropriate to approach miR-182-5p/Nogo-A co-expression studies due to Nogo-A protein expression levels (Fig. 2A) and endogenous expression of miR-182-5p (Fig. 2B). Furthermore, these cell lines have been extensively used to study neuronal differentiation and axonal growth (Tremblay *et al.,* 2010; Watanabe *et al.,* 1994). However, Nogo-A protein is undetected in SH-SY5Y cell line extracts. Thus, we choose Neuro-2a and C6 cell lines to study Nogo-A regulation by miR-182-5p mimic *in vitro*.

**Figure 2.**
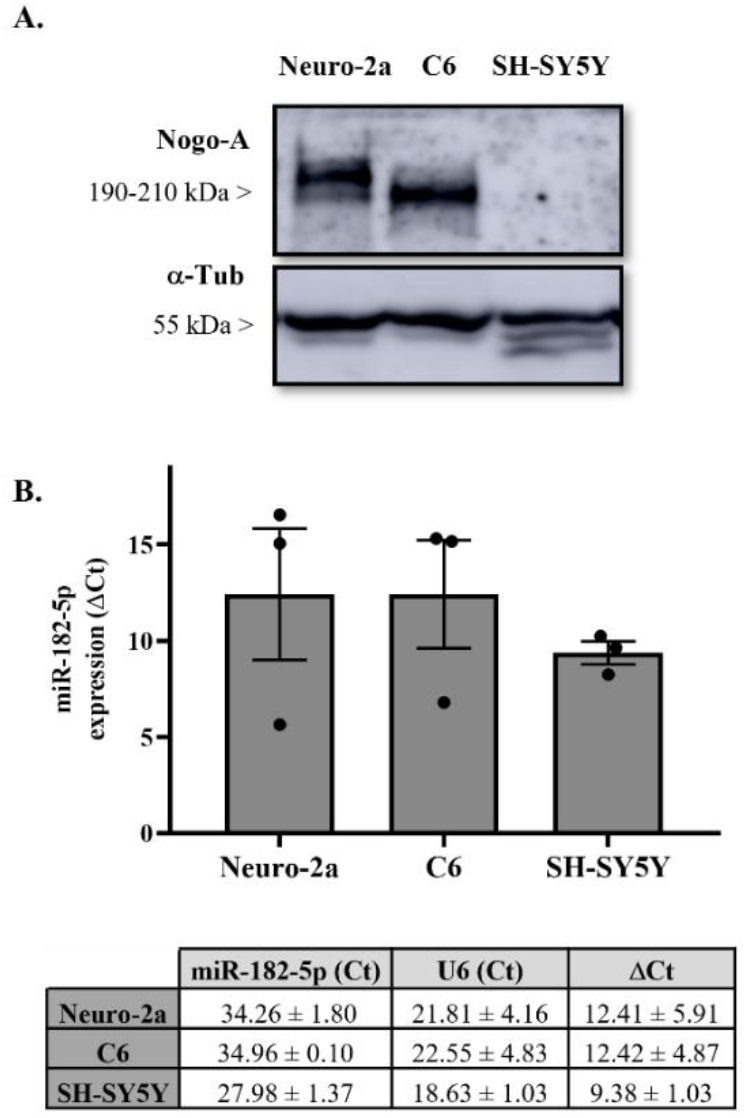
Endogenous expression levels of miR-182-5p and Nogo-A protein in different cell lines. **A)** Representative immunoblot of Nogo-A and load control α-tubulin protein expression in different cell lines lysates of three independent experiments. **B)** RT-qPCR showing relative miR-182-5p expression in total RNA samples isolated from different cell lines. Expression of miR-182-5p from each sample was normalized to the corresponding expression of the control gene snoRNA U6 (ΔCt= Ct_miR182_ – Ct_U6_). The bar graph and table represent the mean ± SEM data of Ct and ΔCt from three independent experiments.

### 3.3. MiR-182-5p targets the mouse Nogo-A 3’UTR and downregulates its protein expression

To confirm the bioinformatics results that miR-182-5p directly targets Nogo-A, Neuro-2a cells were co-transfected with i) the luciferase reporter plasmids containing either the wild-type (pmiRGLO-3’UTR-Nogo-A-wt) or mutant (pmiRGLO-3’UTR-Nogo-A-mut) 3’UTR-Nogo-A and ii) the miR-182-5p or Neg. Ctrl mimics. Co-transfection of pmiRGLO plasmid without 3’UTR insert (pmiRGLO) was used as endogenous control.

The luciferase activity of the pmiRGLO plasmid in presence of miR-182-5p mimic was evaluated to rule out any effect of the microRNA on the plasmid expression, and no significant effects are detected (Fig. 3A) (109.1 ± 21.71% vs. 100% empty pmiRGLO without mimic co-transfection; two-tailed paired t-test, T_4_= 0.4203, n.s. p=0.6958).

**Figure 3.**
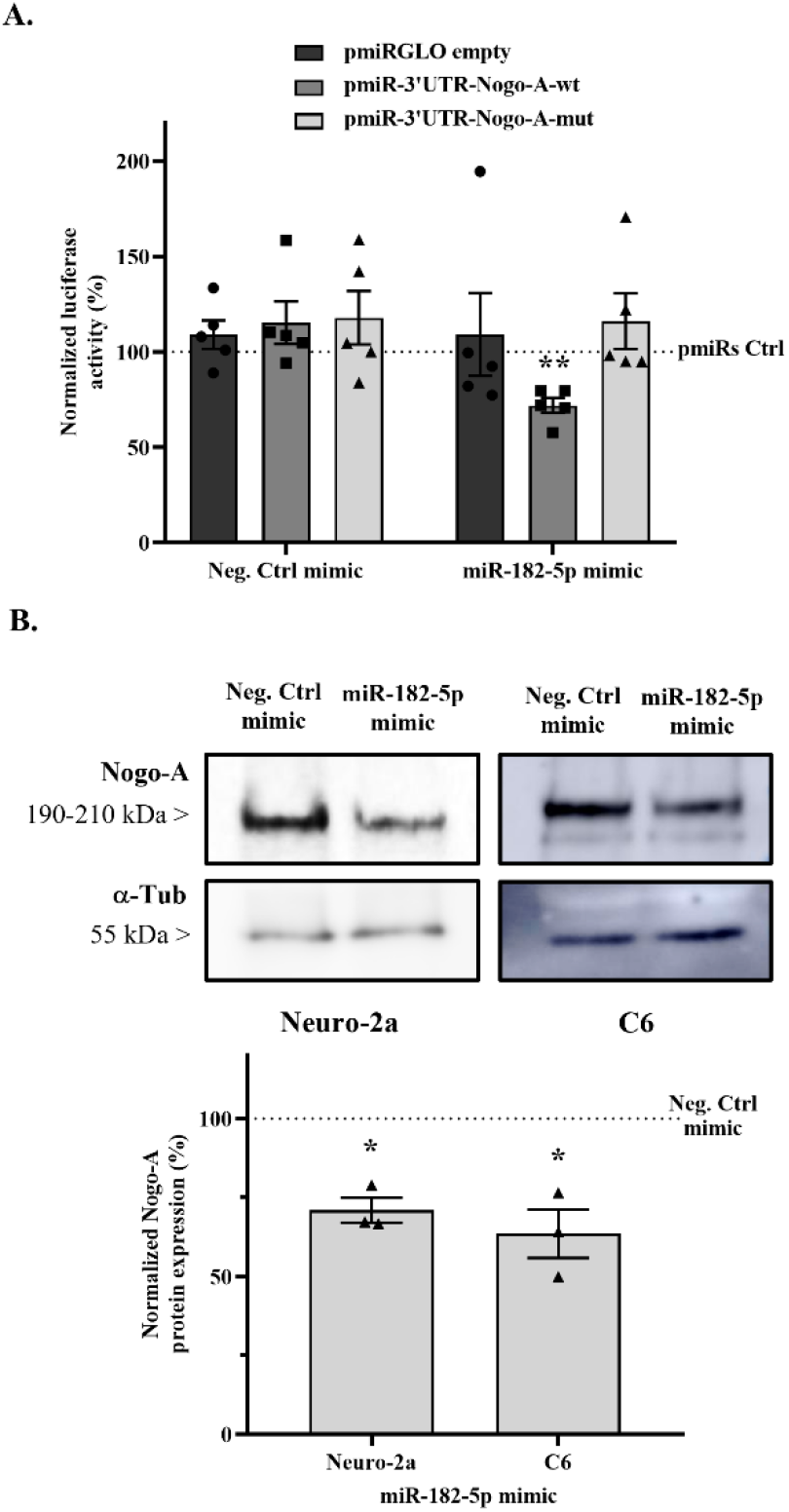
MiR-182-5p mimic regulates Nogo-A expression. **A)** Luciferase reporter activity of either miR-182-5p or Neg. Ctrl mimics on pmiRGLO-3’UTR-Nogo-A-wt, pmiRGLO-3’UTR-Nogo-A-mut, and empty pmiRGLO. Bar graphs show mean ± SEM of the firefly/Renilla emission ratio normalized to their corresponding pmiRGLO without mimic co-transfection (pmiRs Ctrl) in Neuro-2a cells (two-tailed paired t-test, T_4_= 4.789, p=0.0087, n=5 independent experiments). ** indicates a p-value < 0.01. **B)** Representative immunoblot of Nogo-A and the load control α-tubulin protein expression in Neuro-2a cells and C6 cells transfected either with miR-182-5p or Neg. Ctrl mimics. Bar graphs show the mean ± SEM of three independent experiments; Nogo-A expression was normalized by the respective α-tubulin expression taking Neg. Ctrl values as control levels. MiR-182-5p mimic transfection showed statistical reduction in Neuro-2a cell line (two-tailed paired t-test, T_2_=7.291; p= 0.0183) and C6 cell line (two-tailed paired t-test, T_2_=4.755; p= 0.0415). * indicates a p-value < 0.05.

In addition, Neg. Ctrl mimic co-transfection does not affect firefly/Renilla emission ratio of pmiRGLO-3’UTR-Nogo-A-wt (115.24 ± 11.17% vs. 100% pmiRGLO-3′UTR-Nogo-A-wt without mimic co-transfection; two-tailed paired t-test, T^4^=1.365, n.s. p=0.2441) (Fig.3A). However, co-transfection with miR-182-5p mimic causes a significant reduction of a 43.33% of emission ratio compared to co-transfection with the Neg. Ctrl mimic (71.87 ± 4.01% vs. 115.24 ± 11.17% Neg. Ctrl mimic; two-tailed paired t-test, T^4^= 4.789, p=0.0087).

Then, the effects of both mimics on pmiRGLO-3′UTR-Nogo-A-mut plasmid were evaluated, using a mutation in the predicted miR-182-5p binding site, to validate that the effect of miR-182-5p reducing emission ratio is due specifically to the interaction with its binding site. Co-transfection with Neg. Ctrl mimic (117.8 ± 14.01% vs. 100% pmiRGLO-3′UTR-Nogo-A-mut; two-tailed paired t-test, T_4_= 1.273, n.s. p=0.2719) or miR-182-5p mimic (116.1 ± 14.52% vs. 100% pmiRGLO-3′UTR-Nogo-A-mut; two-tailed paired t-test, T_4_= 1.107, n.s. p=0.3304) has no effect on luciferase emission ratio (Fig. 3A).

Finally, to evaluate the modulation of Nogo-A protein expression levels by miR-182-5p, Neuro-2a and C6 cells were transfected with either miR-182-5p or Neg. Ctrl mimics for 24 hours and protein expression levels were detected by immunoblot assay. Transfection with miR-182-5p mimic significantly downregulates the endogenous Nogo-A protein levels compared to Neg. Ctrl mimic transfection in both Neuro-2a cell line (70.98 ± 3.98% vs. 100% Neg. Ctrl mimic; two-tailed paired t-test, T_2_= 7.291, p=0.0183) and C6 cell line (63.51 ± 7.67% vs. 100% Neg. Ctrl mimic; two-tailed paired t-test, T_2_= 4.755, p=0.0415) (Fig. 3B).

### 3.4. Nogo-A downregulation by miR-182-5p mimic promotes neurite outgrowth of rat primary hippocampal neurons

Oligodendrocytic Nogo-A binds to its neuronal receptor (NgR1) and co-receptors (LINGO-1 and p75NTR or TROY) producing the inhibition of neurite outgrowth of the neurons (Fournier *et al.,* 2001; Aloy *et al.,* 2006; Zemmar *et al.,* 2018). To determine the biological effects of the downregulation of Nogo-A by miR-182-5p transfection on neurite outgrowth, we performed functional analyses based on the co-culture of rat primary hippocampal neurons with C6 cells transfected with either miR-182-5p or Neg. Ctrl mimics.

Differences in cell density were observed in the transfected cultures. Transfection of both mimics significantly reduces the C6 cell density in the co-cultures compared to non-transfected C6 cell co-culture (Ctrl co-culture) (two-way ANOVA, F_2,12_=25.52, p<0.0001; Tukey’s multiple comparisons posthoc test, p<0.0001), although hippocampal neurons densities are not significantly changed (Fig. 4B).

**Figure 4.**
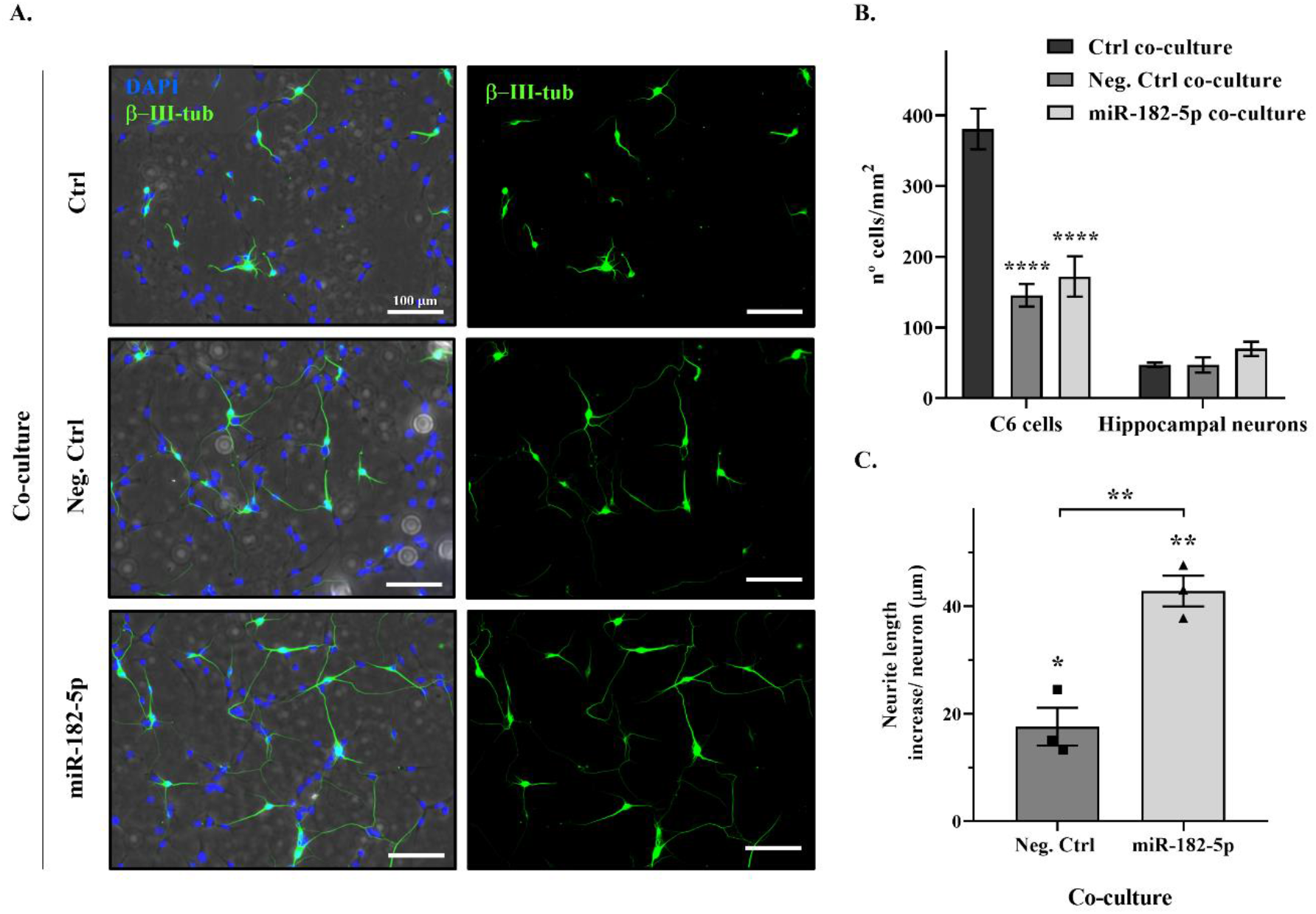
Nogo-A downregulation by miR-182-5p mimic promotes neurite outgrowth of primary hippocampal neurons. **A)** Representative phase contrast and epifluorescence images of different co-cultures of C6 cells (non-transfected control cells; Neg. Ctrl mimic transfected cells; miR-182-5p mimic transfected cells; imaged in phase contrast) and rat hippocampal neurons, labeled with the specific neuronal marker β-III tubulin (green) and DAPI (nuclei staining, blue). Bar scale = 100 μm. **B)** Bar graph shows the mean ± SEM of three independent experiments of the evaluation of the number of C6 cells and hippocampal neurons per mm^2^ (density) in the different co-cultures. Mimics transfection showed statistical reduction in the number of C6 cells (Tukey’s multiple comparisons post-hoc test, p<0.0001). **** indicates a p-value < 0.0001. **C)** Bar graph shows the mean ± SEM of three independent experiments of neurite length increase per neuron (μm) of mimics-transfected C6 cell co-cultures versus the non-transfected C6 cell co-culture values (two-tailed paired t-test; Neg. Ctrl co-culture, T_2_=5.041, p=0.0372; miR-182-5p co-culture, T_2_=15.46, p=0.0042) and the comparison between both mimics-transfected C6 cell co-cultures (two-tailed paired t-test; T_2_=17.65, p=0.0032). * indicates a p-value < 0.05 and ** a p-value < 0.01.

Transfection of C6 cells with miR-182-5p mimic increases significantly neurite length of primary hippocampal neurons in ~26 μm per neuron (miR-182-5p co-culture) in comparison with the Neg. Ctrl transfected C6 cells (Neg. Ctrl co-culture) (183.0 ± 20.32 μm vs. 157.7 ± 20.59 μm Neg. Ctrl co-culture; two-tailed paired t-test, T_2_=17.65, p=0.0032) (Fig. 4A and C).

Moreover, transfection of C6 cells with both Neg. Ctrl and miR-182-5p mimics (Neg. Ctrl co-culture and miR-182-5p co-culture) increases neurite length in comparison with the non-transfected C6 cells (Ctrl co-culture). Neg. Ctrl mimic co-culture induced a significant increase of ~17 μm per neuron (157.7 ± 20.59 μm vs. 140.1 ± 17.58 μm Ctrl co-culture; two-tailed paired t-test, T_2_=5.041, p=0.0372) and miR-182-5p mimic co-culture produced an increase of ~42 μm per neuron (183.0 ± 20.32 μm vs. 140.1 ± 17.58 μm Ctrl co-culture; two-tailed paired t-test, T_2_=15.46, p=0.0042).

## 4. Discussion

Amongst many factors, one of the major inhibitory signals of the CNS environment to regrowth is the myelin-associated Nogo pathway, which plays an important role in regeneration (Baumann *et al.,* 2009; Hånell *et al.,* 2010; Hou *et al.,* 2010). In the present study, we describe and validate for the first time Nogo-A post-transcriptional regulation by a miRNA in murine neural cell lines. We demonstrate that miR-182-5p downregulates Nogo-A expression promoting neurite outgrowth of primary hippocampal neurons *in vitro*.

The involvement of Nogo-A in neurodegeneration has been described in diverse CNS diseases such as ocular diseases, multiple sclerosis, Alzheimer’s disease, amyotrophic lateral sclerosis as well as spinal cord injury (SCI) and brain traumatic injury. However, the role of Nogo-A is not restricted to the CNS, Nogo-A also inhibits the spreading and migration of non-neuronal cells such as fibroblasts and vascular endothelial cells (Lingor *et al.,* 2012; Pernet, 2017).

The dysregulation of Nogo-A following CNS injury, in particular SCI, is in line with the expression changes of numerous genes that play vital roles in the pathogenesis of secondary CNS damage or axonal regeneration (Shi *et al.,* 2017; Li *et al.,* 2021). Most of these genes are regulated by the post-transcriptional regulators miRNAs (Nieto-Diaz *et al.,* 2014) which showed an altered expression following injury (Liu *et al.,* 2009; Yunta *et al.,* 2012; Li and Zhou, 2019; Fei *et al.,* 2021; Wang *et al.,* 2019). Evidence has demonstrated both *in vitro* and *in vivo* that these miRNAs have crosstalk with the key genes involving processes of neuronal plasticity, neuronal degeneration, axonal regeneration, remyelination and, glial scar formation after SCI, through translational repression or mRNA degradation (Wu *et al.,* 2012; Higa *et al.,* 2014; Diao *et al.,* 2015; Fiorenza and Barco, 2016). Thus, miR-133b, miR-135a-5p, and miR-29a regulate neurite outgrowth and axon regeneration via targeting RhoA, ROCK1/2, and PTEN genes, respectively. Moreover, the overexpression of these miRNAs contributed to spinal cord regeneration and functional recovery in murine SCI models (Lu *et al.,* 2015; Theis *et al.,* 2017; Yin *et al.,* 2018; Wang *et al.,* 2020). Similarly, miR-182-5p has been involved in the secondary damage of the CNS processes and neuronal regeneration. According to miRNATissueAtlas2 (https://ccb-web.cs.uni-saarland.de/tissueatlas2, Jan. 2022; Keller *et al.,* 2022), miR-182-5p is mainly expressed in blood vessels, epididymis, and CNS, especially in the spinal cord.

Previous analysis from our laboratory (Yunta *et al.* 2012) and others (Liu *et al.,* 2009; Strickland *et al.* 2011; Zhang and Wu, 2018) revealed miR-182-5p as one of the miRNAs most downregulated after injury, in agreement with the recently described time course miR-182-5p expression results in SCI (Fei *et al.,* 2021). In this latter study, the highest downregulation point of miR-182-5p expression is observed at 7 days post injury with an expression recovery at 28 days that interestingly parallels Nogo-A both protein and mRNA expression which rapidly rose to a peak after 7 days, and then gradually declined again after 14 days (Wang *et al.,* 2015). However, it has not been reported to date a validated miRNA targeting Nogo-A. In accordance with our studies, the modulatory role of miR-182-5p on Nogo-A validated by our luciferase and immunoblot results (Fig. 3) could explain this expression changes dynamic following injury. The validity of the miR-182-5p and Nogo-A interaction is supported by the miR-182-5p regulation of another member of the family RTN4, namely Nogo-C, which shares the same 3’UTR with Nogo-A.

Overexpression of miR-182-5p targets the Nogo-C 3’UTR and decreases its protein levels protecting cardiomyocytes from apoptosis and preserving cardiac function after myocardial infarction (Jia *et al.,* 2016). However, single genes may produce a variety of mRNA isoforms by mRNA modification, such as alternative polyadenylation or splicing that could alter the selective recruitment of miRNAs to the 3’UTR (Fernandez-Moya *et al.,* 2017; Mayr, 2016). It has been observed that nearly all genes have multiple alternative polyadenylation signals, located at different positions in the 3′UTR (Le Hir *et al.,* 2001). Thus, we approach the validation of the miR-182-5p as regulator of Nogo-A, since in both mouse and human NOGO gene, it has been described more than one putative poly(A) signal site downstream of the stop codon and the miR-182-5p site is located between these polyadenylation signals. Our bioinformatics analyses confirmed that miR-182-5p binding site on Nogo-A 3’UTR is conserved across different mammalian species, including human. This miR-182-5p binding site in human Nogo-A 3’UTR has been confirmed by STarMir CLIP-based tool providing for the possibility of Nogo-A regulation by miR-182-5p in human cells. Our experimental data from reporter gene regulation and Nogo-A endogenous expression level after overexpression of miR-182-5p in cell cultures validate this microRNA response element in the Nogo-A 3’UTR.

Although the role of miR-182-5p as a regulator of neurite outgrowth has been described in cortical and midbrain neurons through activation of the PTEN/AKT pathway (Wang *et al.,* 2017; Roser *et al.,* 2018), our results could provide a broader implication with regards to the axonal regeneration. Our functional assays (Fig. 4) showed that the downregulation of Nogo-A due to miR-182-5p overexpressed in neural cells eased the neurite outgrowth of primary hippocampal neurons. However, a better understanding of the miR-182-5p regulation on Nogo-A employing gain- and loss-function assays would be needed to establish its role following CNS injury.

Neutralizing Nogo-A by function-blocking antibodies or genetic knockout (KO) has been shown to improve axonal sprouting and regeneration in the injured spinal cord and brain. (Kim *et al.,* 2003; Dimou *et al.,* 2006; Schwab and Strittmatter, 2014) and the clinical potential of anti-Nogo-A antibodies for managing SCI is currently being investigated in two clinical trials (ClinicalTrials.gov | Identifiers: NCT03935321 and NCT03989440). Therefore, Nogo-A downregulation by overexpression of miR-182-5p could be a potential treatment of different diseases and conditions which implicate axonal degeneration.

Our results provide novel information about the regulatory action of miR-182-5p on the Nogo-A axonal regeneration inhibition. We are first to describe and validate Nogo-A regulation by miR-182-5p which targets Nogo-A 3’UTR, downregulates Nogo-A protein expression levels, and promotes neurite outgrowth in murine neural cell lines. Thus, miR-182-5p could be a promising therapeutic tool for diseases or conditions which implicate axonal pathology such as SCI, brain injury, Parkinson’s, or Alzheimer’s diseases, among others.

## Author Contribution

<u>Altea Soto:</u> Conceptualization, Methodology, Investigation, Writing - Original draft, Writing - Review & Editing, Visualization. <u>Manuel Nieto-Díaz:</u> Validation, Formal analysis, Writing - Review & Editing, Supervision. <u>David Reigada, Teresa Muñoz-Galdeano, and M.</u> <u>Asunción Barreda-Manso:</u> Writing-Reviewing and Editing. <u>Rodrigo M. Maza:</u> Conceptualization, Methodology, Validation, Writing - Review & Editing, Supervision.

## Acknowledgements

We thank the Fundación del Hospital Nacional de Parapléjicos para la Investigación y la Integración (FUHNPAIIN) and the animal facility from the Research Unit of the Hospital Nacional de Parapléjicos (Toledo, Spain) for their technical and logistic support.

## References

Aloy, E. M., Weinmann, O., Pot, C., Kasper, H., Dodd, D. A., Rülicke, T., Rossi, F., and Schwab, M. E. (2006). Synaptic destabilization by neuronal Nogo-A. Brain Cell Biol, 35(2–3), 137–157.

Bartel, D. P. (2004). MicroRNAs: genomics, biogenesis, mechanism, and function. Cell, 116(2), 281–297.

Baumann, M. D., Austin, J. W., Fehlings, M. G., and Shoichet, M. S. (2009). A quantitative ELISA for bioactive anti-Nogo-A, a promising regenerative molecule for spinal cord injury repair. Methods, 47(2), 104–108.

Bellon, A., Iyer, A., Bridi, S., Lee, F. C., Ovando-Vazquez, C., Corradi, E., Longhi, S., Rocuzzo, M., Strohbuecker, S., Naik, S., et al. (2017). miR-182 regulates Slit2-mediated axon guidance by modulating the local translation of a specific mRNA. Cell Rep, 18(5), 1171–1186.

Coolen, M., and Bally-Cuif, L. (2009). MicroRNAs in brain development and physiology. Curr Opin Neurobiol, 19(5), 461–470.

Diao, H. J., Low, W. C., Lu, Q. R., and Chew, S. Y. (2015). Topographical effects on fiber-mediated microRNA delivery to control oligodendroglial precursor cells development. Biomaterials, 70, 105–114.

Dimou, L., Schnell, L., Montani, L., Duncan, C., Simonen, M., Schneider, R., Liebscher, T., Gullo, M., and Schwab, M.E. (2006). Nogo-A-deficient mice reveal strain-dependent differences in axonal regeneration. J Neurosci, 26: 5591–5603.

Fei, M., Li, Z., Cao, Y., Jiang, C., Lin, H., and Chen, Z. (2021). MicroRNA-182 improves spinal cord injury in mice by modulating apoptosis and the inflammatory response via IKKβ/NF-κB. Lab Invest, 1–16.

Fernández‐Moya, S. M., Ehses, J., and Kiebler, M. A. (2017). The alternative life of RNA—sequencing meets single molecule approaches. FEBS lett, 591(11), 1455–1470.

Fiorenza, A., and Barco, A. (2016). Role of Dicer and the miRNA system in neuronal plasticity and brain function. Neurobiol Learn Mem, 135, 3–12.

Fournier, A. E., GrandPre, T., and Strittmatter, S. M. (2001). Identification of a receptor mediating Nogo-66 inhibition of axonal regeneration. Nature, 409(6818), 341–346.

Gu, X., Meng, S., Liu, S., Jia, C., Fang, Y., Li, S., Fu, C., Song, Q., Lin, L., and Wang, X. (2014). miR-124 represses ROCK1 expression to promote neurite elongation through activation of the PI3K/Akt signal pathway. J Mol Neurosci, 52(1), 156–165.

Hånell, A., Clausen, F., Björk, M., Jansson, K., Philipson, O., Nilsson, L. N., Hillered, L., Weinreb, P.H., Lee, D., Mclntosh, T.K., Gimbel, D.A., Strittmatter, M., and Marklund, N. (2010). Genetic deletion and pharmacological inhibition of Nogo-66 receptor impairs cognitive outcome after traumatic brain injury in mice. J Neurotraum, 27(7), 1297–1309.

Higa, G. S. V., de Sousa, E., Walter, L. T., Kinjo, E. R., Resende, R. R., and Kihara, A. H. (2014). MicroRNAs in neuronal communication. Mol Neurobiol, 49(3), 1309–1326.

Hou, T., Shi, Y., Cheng, S., Yang, X., Li, L., and Xiao, C. (2010). Nogo-A expresses on neural stem cell surface. Int J Neurosci, 120(3), 201–205.

Huber, A. B., Weinmann, O., Brösamle, C., Oertle, T., and Schwab, M. E. (2002). Patterns of Nogo mRNA and protein expression in the developing and adult rat and after CNS lesions. J Neurosci, 22(9), 3553–3567.

Hunt, D., Coffin, R. S., Prinjha, R. K., Campbell, G., and Anderson, P. N. (2003). Nogo-A expression in the intact and injured nervous system. Mol Cell Neurosci, 24(4), 1083–1102.

Jia, S., Qiao, X., Ye, J., Fang, X., Xu, C., Cao, Y., and Zheng, M. (2016). Nogo-C regulates cardiomyocyte apoptosis during mouse myocardial infarction. Cell Death Dis, 7(10), e2432–e2432.

Josephson, A., Widenfalk, J., Widmer, H. W., Olson, L., and Spenger, C. (2001). NOGO mRNA expression in adult and fetal human and rat nervous tissue and in weight drop injury. Exp Neurol, 169(2), 319–328.

Keller, A., Gröger, L., Tschernig, T., Solomon, J., Laham, O., Schaum, N., Viktoria W., Kern, F., Schmartz, G. P., Li, Y., et al. (2022). miRNATissueAtlas2: an update to the human miRNA tissue atlas. Nucleic acids Res, 50(D1), D211–D221.

Kim, J.E., Li, S., GrandPre, T., Qiu, D., and Strittmatter, S.M. (2003). Axon regeneration in young adult mice lacking Nogo-A/B. Neuron, 38: 187–199.

Kocerha, J., Kauppinen, S., and Wahlestedt, C. (2009). microRNAs in CNS disorders. Neuromol Med, 11(3), 162–172.

Le Hir, H., Gatfield, D., Izaurralde, E., & Moore, M. J. (2001). The exon–exon junction complex provides a binding platform for factors involved in mRNA export and nonsense‐mediated mRNA decay. EMBO J, 20(17), 4987–4997.

Li, F., and Zhou, M. W. (2019). MicroRNAs in contusion spinal cord injury: pathophysiology and clinical utility. Acta Neurol Belg, 119(1), 21–27.

Li, P., Jia, Y., Tang, W., Cui, Q., Liu, M., and Jiang, J. (2021). Roles of non-coding RNAs in central nervous system axon regeneration. Front Neurosci-Switz, 15.

Lingor, P., Koch, J. C., Tönges, L., and Bähr, M. (2012). Axonal degeneration as a therapeutic target in the CNS. Cell Tissue Res, 349(1), 289–311.

Liu, N. K., Wang, X. F., Lu, Q. B., and Xu, X. M. (2009). Altered microRNA expression following traumatic spinal cord injury. Exp Neurol, 219(2), 424–429.

Livak, K. J., and Schmittgen, T. D. (2001). Analysis of relative gene expression data using real-time quantitative PCR and the 2− ΔΔCT method. Methods, 25(4), 402–408.

Lu, X. C., Zheng, J. Y., Tang, L. J., Huang, B. S., Li, K., Tao, Y., Yu, W., Zhu, R.L., Li, S., and Li, L. X. (2015). MiR-133b Promotes neurite outgrowth by targeting RhoA expression. Cell Physiol Biochem, 35(1), 246–258.

Mayr, C. (2016). Evolution and biological roles of alternative 3′ UTRs. Trends Cell Biol, 26(3), 227–237.

Montani, L., Gerrits, B., Gehrig, P., Kempf, A., Dimou, L., Wollscheid, B., and Schwab, M. E. (2009). Neuronal Nogo-A modulates growth cone motility via Rho-GTP/LIMK1/cofilin in the unlesioned adult nervous system. J Biol Chem, 284(16), 10793–10807.

Moore, M. J., Scheel, T. K., Luna, J. M., Park, C. Y., Fak, J. J., Nishiuchi, E., Rice, C.M., and Darnell, R. B. (2015). miRNA–target chimeras reveal miRNA 3′-end pairing as a major determinant of Argonaute target specificity. Nat Commun, 6(1), 1–17.

Nieto-Diaz, M., Esteban, F. J., Reigada, D., Muñoz-Galdeano, T., Yunta, M., Caballero-López, M., Navarro-Ruiz, R., del Águila, A., and Maza, R. M. (2014). MicroRNA dysregulation in spinal cord injury: causes, consequences and therapeutics. Front Cell Neurosci, 8, 53.

Oertle, T., and Schwab, M. E. (2003). Nogo and its paRTNers. Trends Cell Biol, 13(4), 187–194.

Oertle, T., Huber, C., van der Putten, H., and Schwab, M. E. (2003). Genomic structure and functional characterisation of the promoters of human and mouse nogo/rtn4. J Mol Biol, 325(2), 299–323.

Pernet, V. (2017). Nogo-A in the visual system development and in ocular diseases. BBA-Mol Basis Dis, 1863(6), 1300–1311.

Rennie, W., Liu, C., Carmack, C. S., Wolenc, A., Kanoria, S., Lu, J., Long, D., and Ding, Y. (2014). STarMir: a web server for prediction of microRNA binding sites. Nucleic acids Res, 42(W1), W114–W118.

Rønn, L. C., Ralets, I., Hartz, B. P., Bech, M., Berezin, A., Berezin, V., Møller, A., and Bock, E. (2000). A simple procedure for quantification of neurite outgrowth based on stereological principles. J Neurosci Meth, 100(1–2), 25–32.

Roser, A. E., Gomes, L. C., Halder, R., Jain, G., Maass, F., Tönges, L., Tatenhorst, L., Bähr, M., Fischer, A., and Lingor, P. (2018). miR-182-5p and miR-183-5p act as GDNF mimics in dopaminergic midbrain neurons. Mol Ther-Nucl Acids, 11, 9–22.

Schwab, M. E. (2010). Functions of Nogo proteins and their receptors in the nervous system. Nat Rev Neurosci, 11(12), 799–811.

Schwab, M. E., and Strittmatter, S. M. (2014). Nogo limits neural plasticity and recovery from injury. Curr Opin Neurobiol, 27, 53–60.

Shi, L. L., Zhang, N., Xie, X. M., Chen, Y. J., Wang, R., Shen, L., Zhou, J. G., and Lü, H. Z. (2017). Transcriptome profile of rat genes in injured spinal cord at different stages by RNA-sequencing. BMC genomics, 18(1), 1–14.

Strickland, E. R., Hook, M. A., Balaraman, S., Huie, J. R., Grau, J. W., and Miranda, R. C. (2011). MicroRNA dysregulation following spinal cord contusion: implications for neural plasticity and repair. Neuroscience, 186, 146–160.

Theis, T., Yoo, M., Park, C. S., Chen, J., Kügler, S., Gibbs, K. M., and Schachner, M. (2017). Lentiviral delivery of miR-133b improves functional recovery after spinal cord injury in mice. Mol Neurobiol, 54(6), 4659–4671.

Tremblay, R. G., Sikorska, M., Sandhu, J. K., Lanthier, P., Ribecco-Lutkiewicz, M., and Bani-Yaghoub, M. (2010). Differentiation of mouse Neuro 2A cells into dopamine neurons. J Neurosci Meth, 186(1), 60–67.

Wang, F., Zhong, S., Zhang, H., Zhang, W., Zhang, H., Wu, X., and Chen, B. (2015). Prognostic value of MicroRNA-182 in cancers: A meta-analysis. Dis markers, 2015.

Wang, J. W., Yang, J. F., Ma, Y., Hua, Z., Guo, Y., Gu, X. L., and Zhang, Y. F. (2015). Nogo-A expression dynamically varies after spinal cord injury. Neural Regen Res, 10(2), 225.

Wang, N., Yang, Y., Pang, M., Du, C., Chen, Y., Li, S., Tian, Z., Feng, F., Wang, Y., Chen, Z., et al. (2020). MicroRNA-135a-5p Promotes the Functional Recovery of Spinal Cord Injury by Targeting SP1 and ROCK. Mol Ther-Nucl Acids, 22, 1063–1077.

Wang, W. M., Lu, G., Su, X. W., Lyu, H., and Poon, W. S. (2017). MicroRNA-182 regulates neurite outgrowth involving the PTEN/AKT pathway. Front Cell Neurosci, 11, 96.

Wang, W., Su, Y., Tang, S., Li, H., Xie, W., Chen, J., Shen, L., Pan, X., and Ning, B. (2019). Identification of noncoding RNA expression profiles and regulatory interaction networks following traumatic spinal cord injury by sequence analysis. Aging (Albany NY), 11(8), 2352.

Watanabe, E., Hosokawa, H., Kobayashi, H., and Murakami, F. (1994). Low Density, but not High Density, C6 Glioma Cells Support Dorsal Root Ganglion and Sympathetic Ganglion Neurite Growth. Eur J Neurosci, 6(8), 1354–1361.

Wei, Q., Lei, R., and Hu, G. (2015). Roles of miR‐182 in sensory organ development and cancer. Thorac Cancer, 6(1), 2–9.

Wu, D., and Murashov, A. K. (2013). MicroRNA-431 regulates axon regeneration in mature sensory neurons by targeting the Wnt antagonist Kremen1. Front Mol Neurosci, 6, 35.

Wu, D., Raafat, A., Pak, E., Clemens, S., and Murashov, A. K. (2012). Dicer-microRNA pathway is critical for peripheral nerve regeneration and functional recovery in vivo and regenerative axonogenesis in vitro. Exp Neurol, 233(1), 555–565.

Yin, H., Shen, L., Xu, C., and Liu, J. (2018). Lentivirus-mediated overexpression of miR-29a promotes axonal regeneration and functional recovery in experimental spinal cord injury via PI3K/Akt/mTOR pathway. Neurochem Res, 43(11), 2038–2046.

Yu, B., Qian, T., Wang, Y., Zhou, S., Ding, G., Ding, F., and Gu, X. (2012). miR-182 inhibits Schwann cell proliferation and migration by targeting FGF9 and NTM, respectively at an early stage following sciatic nerve injury. Nucleic acids Res, 40(20), 10356–10365.

Yunta, M., Nieto-Diaz, M., Esteban, F. J., Caballero-López, M., Navarro-Ruíz, R., Reigada, D., Pita-Thomas, D.W., del Águila, A., Muñoz-Galdeano, T., and Maza, R. M. (2012). MicroRNA dysregulation in the spinal cord following traumatic injury. PloS one, 7(4), e34534.

Zemmar, A., Chen, C. C., Weinmann, O., Kast, B., Vajda, F., Bozeman, J., Isaad, N., and Schwab, M. E. (2018). Oligodendrocyte-and neuron-specific nogo-a restrict dendritic branching and spine density in the adult mouse motor cortex. Cereb Cortex, 28(6), 2109–2117.

Zhang, J., and Wu, Y. (2018). microRNA-182-5p alleviates spinal cord injury by inhibiting inflammation and apoptosis through modulating the TLR4/NF-κB pathway. Int J Clin Exp Patho, 11(6), 2948.

Zhou, S., Shen, D., Wang, Y., Gong, L., Tang, X., Yu, B., Gu, X., and Ding, F. (2012). microRNA-222 targeting PTEN promotes neurite outgrowth from adult dorsal root ganglion neurons following sciatic nerve transection. PloS one, 7(9).

